# Non-Thermal Plasma Activated Water is an Effective Nitrogen Fertilizer Alternative for *Arabidopsis thaliana*

**DOI:** 10.1101/2025.06.12.659237

**Authors:** Jonathan Kizer, Conner Robinson, Ta’Kia Lucas, Steven Shannon, Ricardo Hernández, Katharina Stapelmann, Marcela Rojas-Pierce

## Abstract

Nitrogen (N) fixation with non-thermal plasmas has been proposed as a sustainable alternative to meet growing N fertilizer demands for agriculture. This technology generates Plasma Activated Water (PAW) with a range of chemical compositions, including different concentrations of nitrate (NO₃⁻) and hydrogen peroxide (H_2_O_2_), among other compounds. Potential use of PAW as an effective crop fertilizer necessitates a robust understanding of the underlying biology of the plant, which is not yet available. The lack of a unified standard in PAW production and the varying chemical make-up that results from different devices and protocols hampers comparative studies and adoption of this technology. The objective of this study was to compare the efficacy of two PAW solutions with differing concentrations of H_2_O_2_ produced from a Radio Frequency (RF) glow discharge plasma source. The effect of these solutions on plant growth, ROS accumulation, gene expression and heat stress response were compared to N-equivalent controls in the model plant Arabidopsis to assess their potential as an alternative N fertilizer. While PAW solutions lacking detectable H_2_O_2_ enhanced seedling growth, those containing approximately 0.3 µM of H_2_O_2_ did not. ROS accumulation in root tissues was similar between PAW and chemically equivalent solutions, suggesting H_2_O_2_ is the primary ROS present in the PAW at the time of treatment. Gene expression studies showed induction of genes involved in N uptake and assimilation in PAW-treated seedlings. Pre-treatment of seedlings with PAW solutions containing H_2_O_2_ improved root growth under heat stress which indicates that this treatment may induce plant stress response pathways. Finally, mature plants showed similar growth when fertilized with PAW lacking H_2_O_2_ or NO_3_^-^ control regimes for over 5 weeks indicating equivalency in chemical composition, plant nutrient uptake and utilization. Overall, these results demonstrate that PAW is an effective alternative to NO_3_^-^ fertilizers for plant cultivation but the levels of H_2_O_2_ need to be carefully controlled.

## Introduction

The demand for nitrogen (N) fertilizer is concomitant to the rising global demands for food (Lassaletta et al., 2014). In the last 50 years, synthetic fertilizers have made up the largest portion of N fertilizers utilized in global agriculture (Lassaletta et al., 2014). The most used method to fix N for fertilizers is the Haber-Bosch process which is energy intensive, reliant on fossil-fuels, and contributes 2.9 tons of atmospheric CO_2_ globally annually (M. Wang et al., 2021). Alternative N-fixation strategies are being investigated to enhance the sustainability of agricultural practices, while meeting growing fertilizer demands (Chen et al., 2018). Non-thermal plasmas (NTP) can be utilized to fix gaseous N_2_ from the atmosphere into water in a plant-available N form providing a potential alternative method of N fertilizer production. Reactive oxygen and nitrogen species (RONS) can also be produced during plasma treatment of water where specific parameters of the treatment process can affect the resulting chemistry and concentration of RONS (Bae et al., 2024; Lietz & Kushner, 2016; Robinson & Stapelmann, 2024). These RONS are also relevant to plant health. The resulting solution is referred to as Plasma-Activated Water (PAW) or Plasma-Treated Water (PTW).

Plant health is dependent on the availability of nutrients in their environment. Chief among these nutrients are the macronutrients nitrogen (N), phosphorus (P), and potassium (K), with N being vital for plant productivity (Zhang et al., 2022). Plants preferentially take up NO_3_^−^ and NH_4_^+^ from the soil, with NO_3_^−^ being a major source for N for many plants (Bloom, 2015). Roots primarily take up NO_3_^−^ via high- and low-affinity transporters located in the plasma membrane, including those from the NRT1 and NRT2 protein families (Krapp, 2015). Once in the cytosol, NO_3_^−^ is first reduced to NO_2_^−^ by the enzyme nitrate reductase, and NO_2_^-^ is later transported to the chloroplast to be further reduced to NH_4_^+^ by nitrite reductase. NH_4_^+^ can then be incorporated into organic N compounds such as amino acids (Krapp, 2015). NH_4_^+^ can also be taken up directly from the soil via high- and low-affinity transporters. Ammonia Transporter (AMT) family transporter proteins are primarily responsible for NH_4_^+^ uptake in the root (Gansel et al., 2001). The expression of AMT transporters, is highly regulated and controlled by factors such as light, external pH, and internal NH_4_^+^ levels (Loqué & von Wirén, 2004).

N uptake and assimilation are regulated by N availability in the immediate soil environment. NO_3_^−^ functions as a signal molecule that ultimately controls the response to N availability. The perception of this signal leads to downstream responses involving transcriptional, post-transcriptional, and post-translational control (Bouguyon et al., 2012). Some members of the NRT1 and NRT2 family of membrane transporters have additional functions as NO_3_^-^ receptors, with NRT1.1 likely playing this role in root cell plasma membranes (Bouguyon et al., 2015; Maghiaoui et al., 2020).

NO_3_^-^ can control development and morphology of both belowground and aboveground organs (Bouguyon et al., 2015). When nitrate concentration is low (< 1 mM in *Arabidopsis thaliana*), there is overall more elongation in both primary and secondary roots, and this is described as a “foraging” response. This response increases the root surface area and enhances uptake of NO_3_^-^ from surrounding soil. In contrast, plants growing in high NO_3_^-^ conditions (> 10 mM in *A. thaliana*) have reduced root elongation, and lateral roots become repressed at the seedling stage (Bouguyon et al., 2012). These morphological responses are integrated alongside phytohormone response. N-driven lateral root initiation is regulated by the crosstalk between NO_3_^-^ and auxin pathways (Maghiaoui et al., 2020). Moreover, the NO_3_^-^ transceptor NRT1.1 likely has auxin transport capability, in which it basipetally transports auxin from the lateral root primordia, preventing accumulation and subsequent elongation of the lateral roots in response to ample NO_3_^-^ (Bouguyon et al., 2015; Krouk et al., 2010). Overall, the response to NO_3_^-^ is highly regulated and dependent on the availability of the nutrient in the root environment.

Reactive oxygen species (ROS) are naturally produced in chloroplast and mitochondria in photosynthetic tissues and mainly mitochondria in non-photosynthetic tissues. In plant cells, ROS are produced as a byproduct due to natural inefficiencies of the electron transport chain. Peroxisomes, glyoxysomes, and the apoplast are also sites of ROS production with the latter occurring via cell wall peroxidases and plasma membrane NADPH oxidases. While prevalent in the cell, ROS can be detrimental to plant growth. ROS accumulation ultimately can lead to cell dysfunction or death by distorting redox homeostasis and potentially damaging proteins, cellular membranes, and nucleic acids (Singh et al., 2016). Plant cells maintain several scavenging mechanisms to curtail excessive damage from excessive ROS, including enzymes that catalyze the breakdown of the reactive species to more stable species. These include Superoxide Dismutases (SOD) which catalyze the conversion of superoxide (O_2_^-^) to H_2_O_2_ (Kliebenstein et al., 1998; Pilon et al., 2011) and Catalases (CAT) which catalyzes the conversion of H_2_O_2_ into H_2_O (Bueso et al., 2007; Dvořák et al., 2021; Tuzet et al., 2018). Additionally, there is ample evidence that ROS are required in biological systems (Singh et al., 2016; Tsukagoshi, 2016). For example, hydrogen peroxide (H_2_O_2_) and superoxide (O_2_^-^) modulate plant growth, development (Chapman et al., 2019; Dunand et al., 2007) and stress responses (Tsukagoshi et al., 2010).

Both ROS and reactive nitrogen species (RONS) can function as signaling molecules and lead to acclimation to varying abiotic stresses (Chapman et al., 2019; Yao et al., 2017). Abiotic stressors may result in a build-up of RONS beyond the scavenging ability of antioxidants, and RONS accumulation can elicit responses at the cellular and whole-plant levels (Gilroy et al., 2016; Mittler & Blumwald, 2015). Incidentally, introduction to RONS prior to a stress event can induce tolerance to that stress including nutrient and water deficit in a response referred to as “priming” (Rodríguez-Ruiz et al., 2019; Savvides et al., 2016; Zhu et al., 2019). Altogether, these complex responses to exogenous and endogenous RONS may be leveraged to improve growing practices in stressful environments (Chen et al., 2023; Wang et al., 2023).

N fixation via non-thermal plasma represents an alternative to supplement growing global demands for NO_3_^-^ (Chen et al., 2018; Lamichhane et al., 2021; Ndiffo Yemeli et al., 2021; Ran et al., 2024; Rathore et al., 2022; Y. Wang et al., 2021). Studies of the effects of PAW on varying crop species have begun to describe the potential of plasma agriculture (Adhikari et al., 2019; Kučerová et al., 2021; Ndiffo Yemeli et al., 2021; Škarpa et al., 2020; Zambon et al., 2020). From these studies, several crops showed increased growth when treated with PAW, with cereal crops notably showing significant height differences compared to untreated controls (Ndiffo Yemeli et al., 2021; Škarpa et al., 2020). However, while several of these studies hypothesized that plant growth differences are due to the ROS present in PAW, few studies have delved deeper into the underlying biology. Studies in model plant systems do exist with some focusing more on direct-plasma treatment of seedlings (Cui et al., 2021; Cui et al., 2022). Others in the field have begun specifically investigating the underlying effects of PAW-treatment on the plant response (Cortese et al., 2021; Ka et al., 2021) and how these responses can bolster the efficacy of PAW as a supplemental source of N. However, the field of plasma agriculture has no unified standard in how the impacts of PAW to plants is quantified to determine whether the effects of PAW treatment are applicable to other systems and environments.

Here, we compared the growth of Arabidopsis plants under PAW treatment or equivalent fertilizer regimes using traditional sources of N. We found that PAW treatments with low ROS concentrations resulted in comparable plant growth throughout the plant lifecycle when compared to a simulated NO_3_^-^ fertilizer. Small increases in root growth were observed in PAW-treated plants compared to an equivalent NO_3_^-^ control. We did not find evidence that PAW treatment affected the accumulation of ROS or hormone response markers in plant roots. Enhanced tolerance to heat stress was observed in seedlings pre-treated with H_2_O_2_-containing PAW. Our studies underscore the potential of PAW as an alternative to NO_3_^-^ fertilizers in plants.

## Materials and Methods

### PAW production

Plasma water treatments were conducted using an atmospheric radio frequency (RF) glow discharge plasma, as described in (Byrns et al., 2012). This plasma device was used to surface treat bulk DI water volumes to produce the PAW used in these experiments. An AE OVAtion 35162 RF Generator is used to power the device, and the corresponding software is used to control the delivered power. For all treatments, the delivered power was kept constant at 250 W. Air was flowed down the coaxial electrode, toward the water surface at a rate of ≤ 1 slm (this was controlled with an analog gauge which did not read below this value and would vary depending on the background pressure from the buildings compressed air supply). Treatment parameters varied depending on the desired chemistry, namely the inclusion of H_2_O_2_. To optimize the production of NO_3_^-^, a large external volume of water (≥ 2 L) was circulated through the plasma chamber - which was kept open to improve ventilation - and the distance between the water surface and electrode was set to minimize reflected power (20–40 W reflected at 1.5 cm). Under these conditions, the plasma was consistently able to produce aqueous NO_3_^-^ at a rate of 2 mg/min and treatment times were adjusted accordingly to achieve the desired concentration for the target volume. For H_2_O_2_ production, the plasma chamber was sealed with a smaller, stagnant water volume (450 mL), and the gap distance was reduced to roughly 0.75 cm (100–200 W reflected). This would result in significant evaporation, lowering the water surface and increasing the gap distance as the treatment progressed. The reflected power corresponded closely with the gap distance (when other parameters are kept constant) and was used to determine when the gap distance had changed significantly. The water volume was placed upon a lab jack within the chamber, and when the reflected power dropped below the desired range, the jack would be raised by hand while the treatment was ongoing. These treatments lasted 45 minutes, with 250 ml of DI water remaining by the end. This PAW typically contained 50 mg/L of H_2_O_2_ and 50–100 mg/L of nitrate NO_3_^-^; however, this was far less consistent than the NO_3_^-^ focused treatments. Treatments were repeated with fresh DI water until the amount of aqueous H_2_O_2_ necessary to achieve the desired concentration for the target volume was produced. The NO_3_^-^ only and NO_3_^-^ + H_2_O_2_ PAW were initially kept separate, their chemical composition measured colorimetrically and then were combined with one another and/or untreated DI water to obtain the final desired PAW volumes and chemistries. The final PAWs were also tested colorimetrically to confirm their composition. NO_3_^-^, NO_2_^-^, H_2_O_2_, and NH_4_^+^ levels were tested for each step in triplicate using the commercially available Supelco test kits: 1.09713, 1.14776, 1.18789 and 1.14752, respectively. Absorbance values were obtained using an UV-VIS-NIR light source (Ocean Optics DH-2000-BAL) in conjunction with a spectrometer (Ocean Optics QE65 Pro) and a cuvette holder (Ocean Optics CUV-UV). These absorbances were converted to concentrations using stock solution based standard curves prepared in advance. PAW was neutralized with 1M KOH solution to increase the pH to 5.7, a plant-viable pH. Neutralization maintained stability of the solution for storage. PAW was stored in the dark at room temperature for up to 2 weeks before use.

### Plant Material and Growth Conditions

*Arabidopsis thaliana* ecotype Columbia 0 (Col-0) was used for all experiments. Arabidopsis lines expressing the hormone response markers DR5::GFP (Ulmasov et al., 1997) or TCSn::GFP (Zürcher et al., 2013) were obtained from the Arabidopsis Biological Resource Center (ABRC). The EBS:Ypet marker line was previously described (Fernandez-Moreno & Stepanova, 2020). Seeds were surface sterilized with 95% ethanol followed by a solution containing 20% commercial bleach and 0.1 % Tween 20 (VWR, MFCD00165986). Seeds were rinsed 2-3 times with sterile diH_2_O and then stored at 4 °C for 4 days in the dark. Seeds were then plated onto Arabidopsis Growth Media (AGM) containing 0.5 x MS with MES (Murashige & Skoog, 1962, RPI, M70300), 1% sucrose and 4g/L Gelrite (RPI, G35020). Plates were incubated vertically in a growth chamber with 120 µmol/m^2^/s of PPFD at 22 °C with a 16 h/8 h day/night cycle to promote germination. Plants potted in soil were grown in growing benches with LED grow lights under similar controlled conditions as plated plants. PPFD was provided at 130 µmol/m^2^/s at plant height. Plants grown on soil were rotated to new positions regularly on grow benches to reduce impact of uncontrolled environmental effects.

### PAW and NO_3_^-^ control treatments

AGM with PAW or NO_3_^-^ for seedling treatments was prepared as follows: 1 X (4.3 g/L) MS media without nitrogen (MS-N, Bio-World, 30630200) was adjusted to pH 5.7, supplemented with 8 g/L Gelrite and 2% sucrose and then sterilized by autoclaving (2X AGM-N). Control solutions were prepared in diH_2_O using potassium nitrate (KNO_3_^-^) (Caisson Labs, P012) and 30% (w/w) H_2_O_2_ solution (Sigma-Aldrich, H1009) at the same concentrations of pre-mixed PAW (4.8 mM or 3.5 mM NO_3_^-^ with or without 0.3 mM H_2_O_2_). All treatment solutions including PAW were gently brought up to 60 °C in a water bath and then filter sterilized using a 0.45 µm pore filter inside a laminar flow hood. The 2X AGM-N media was melted in a microwave and cooled to 60 °C just before mixing. Treatment solutions were then mixed with an equal volume of 2X AGM-N media in a pre- heated bottle and the mixture was rapidly poured into square petri dishes in the laminar flow hood.

PAW treatments at the seedling stage were performed by transferring 4-day old seedlings from AGM media to AGM containing PAW or control solutions and incubated for 5-7 days or as indicated. PAW treatments in soil were achieved by irrigation of nutrient solutions as follows. A substrate mixture devoid of N was made to avoid any unspecified fertilizer normally present in commercial soil mixes. The soil mixture contained 45% peat moss (Premier Peat Moss), 35% vermiculite (Sta-Green Vermiculite), and 20% perlite (Aero Soil Perlite) (all measured by volume). We sourced soil components that were not amended with nutrients as per the manufacturer’s label. Pulverized limestone (Gardenlime) was added to adjust pH to ∼6.0. The soil was moistened with deionized H_2_O and distributed into 2-inch insert pots (T.O. Plastics, 2401 Standard). Three-day old seedlings previously germinated on sterile AGM media were transferred to each pot of moistened soil. Seedlings were thinned to 1 per pot 3 days after transfer. All seedlings were watered with 50 ml diH_2_O per pot twice weekly to keep the plants hydrated. 50 ml per pot of 0.25x Hoagland Solution without N (Bio-World, 30630038) was given to all seedlings biweekly by top irrigation. Specific treatments were provided by top irrigation with 50 ml per pot of either PAW, equivalent NO_3_^-^ control solution, or Low N (0.5 mM NO_3_^-^) control solution once per week.

### Plant Growth Measurement

Seedling plates were scanned using a desktop scanner every 2 days and the primary and lateral root lengths were measured using ImageJ software version Fiji (Schindelin et al., 2012). Total root length was calculated by summing the primary and lateral root lengths. Lateral root density was calculated by dividing the number of lateral roots of a given individual by the primary root length.

After 5 weeks of treatment, soil-grown plants were imaged top-down with a 12 MP digital camera positioned 18 cm above the soil. These images were used to calculate rosette area using Fiji. After imaging, soil-grown plants were harvested, and shoots were separated from roots. Aboveground tissue fresh biomass was obtained immediately. Then shoots and cleaned root samples were dried at 175°C for 12 hours to determine dry biomass.

### Heat Stress Experiments

Four-day old seedlings, grown as described above, were transferred onto PAW or control treatment plates and incubated for 1 day at 22°C. Heat stress was achieved by transferring half of the plates to an incubator set at constant temperature of 30°C while keeping identical PPFD and day/night lighting cycles. The remaining plates were kept at 22°C as controls. After 4 days, all plates were scanned with a desktop scanner, and their root lengths were measured using ImageJ. The experiments were replicated 3 times with similar results.

### Fluorescent probe staining, imaging, and quantification

Four-day old seedlings were transferred onto AGM media containing PAW, NO_3_^-^ or NO_3_^-^ combined with H_2_O_2_ (0.15mM or 10mM) and stained 2 minutes, 2 hours or 2 days later with 2’,7’ Dichlorodihydrofluorescin Diacetate (H_2_DCFDA, Sigma-Aldrich, 287810) or Peroxy Orange 1 (PO1, Tocris Bioscience, 49-441-0). H_2_DCFDA and PO1 were dissolved in DMSO as 10 mM stocks. Dyes were diluted to 10 μM (H_2_DCFDA) or 50 μM (PO1) working solutions in 0.5 x AGM-N liquid media and seedlings were stained in the dark for 10 min (H_2_DCFDA) or 30 min (PO1) before one rinse with water and imaging in the microscope. Stained seedlings were imaged on a Zeiss LSM 980 confocal microscope using a 20x objective (Zeiss Plan-Apochromat/ 0.8 N.A.). H_2_DCFDA was excited with a 488 nm laser at 0.25% maximum power and emission was collected at 509-550 nm. PO1 was excited with a 488 nm laser at 0.1% maximum power and emission was collected at 544-695 nm.

### RNA sequence analysis

Three-day old Col-0 Arabidopsis seedlings were transferred onto AGM containing PAW4 or NO_3_^-^ and grown for additional 8 days. The shoot and root tissue of these seedlings were excised, and excess gel media was removed immediately by gentle blotting on low-lint tissue paper. Samples were pooled into four replicates of approximately 20 Arabidopsis seedlings each and placed in 5 volumes of RNAlater stabilization solution (Thermo Fisher Scientific, AM7020) based on tissue mass as specified by the manufacturer. Seedlings spent less than 10 seconds from harvesting to transfer to RNAlater. RNA was then extracted from the tissue using the RNeasy plant mini kit (Qiagen, 74904). Quality of extracted RNA was tested utilizing an Agilent 4200 Tapestation. cDNA library preparation was performed for samples with passing RIN scores using the NEB NEBNext Ultra II Direction RNA kit (polyA enriched) (NEB, E7760S). Next-Generation Sequencing was conducted using an Illumina NovaSeq 6000 with 150-base paired-end reads. The resulting sequencing data was processed and analyzed utilizing the Qiagen CLC Genomics workbench. Expression data for root and shoot data was analyzed for replicates treated with PAW4 against replicates treated with the NO_3_^-^ control. Root and shoot tissues were analyzed separately. Gene ontologies were investigated and plotted utilizing the ShinyGO (version 0.82) bioinformatics tool with a focus on biological processes (Ge et al., 2019).

### Statistical Analysis

Statistical analysis was performed in this study utilizing the R programming language (R-Project). For root length comparisons, a one-way ANOVA with significance at a 0.05 alpha was performed to determine significant differences in length across multiple treatments. A Tukey HSD test was performed as a post-hoc analysis to determine specific relationships between treatments. Tukey HSD was chosen for the test’s effectiveness when performing pairwise comparisons and ability to reduce false positive errors (Agbangba et al., 2024). For experiments comparing the effects of treatment and elevated temperature on root length, a two-way ANOVA was conducted. Comparisons were made across media treatments and temperatures. Tukey HSD was conducted as a pos-hoc analysis.

Analysis of tissue biomass, plant heights, and rosette areas was performed with a Student’s T-test. All data was assumed normal, and significance was denoted at a 0.05 alpha.

## RESULTS

### Optimal PAW treatments promote total root elongation in seedlings outside the effects of NO_3_^-^ and H_2_O_2_

In order to test the effect of multiple PAW chemistries on plant growth, a Radio Frequency (RF) glow discharge plasma source (Byrns et al., 2012; Lindsay et al., 2014) was used to produce PAW with differing concentrations of NO_3_^-^ or H_2_O_2_ (Table 1), and this allowed pairwise comparisons. PAW treatments after incorporation into gel media contained either 1.75 mM (110 ppm, PAW1 and PAW2) or 2.4 mM (150 ppm, PAW4 and PAW5) of NO_3_^-,^ which correspond to NO_3_^-^ sufficient concentrations for Arabidopsis growth (Boer et al., 2020). PAWs differed within each pair by the presence of 0.15 mM H_2_O_2_ such that the effect of this ROS could be assessed systematically. The RF glow discharge plasma source was suitable for producing 2 mg/min NO_3_^-^ in deionized H_2_O and thus was feasible for plant experiments. PAW treatments were applied to 4 day-old Arabidopsis (Col-0) seedlings to circumvent well-documented effects of PAW on seed germination (Bafoil et al., 2019; Cui et al., 2021; Cui et al., 2019; Kučerová et al., 2019). Seedlings were transferred to N-free media supplemented with PAW, diH_2_O (negative control), 1.75 mM NO_3_^-^ or 1.75 mM NO_3_^-^ combined with 0.15 mM H_2_O_2_. Seedling growth was measured after 5 days of treatment, which was 9 days after germination. Plants treated with NO_3_^-^ controls or any PAW solution showed a significant increase in root length compared to diH_2_O controls as expected (Fig. 1A, B). PAW1 and PAW4 treatments, which contain 1.75 and 2.4 mM NO_3_^-^, respectively, but no measurable H_2_O_2_, showed ∼11% increase in total root length compared to the NO_3_^-^ control. Interestingly, PAW2 treatment resulted in a 14% decrease in total root length on average when compared to PAW1, suggesting that H_2_O_2_ in this PAW may dampen root elongation. This effect was also evident in the shorter total and primary roots of the NO_3_^-^ plus H_2_O_2_ controls when compared to NO_3_^-^ alone. PAW5 treatments resulted in similar total root length as the PAW4-treated plants. Measurements of primary root length showed similar results where PAW2 and PAW5 treatments, which contain measurable NO_3_^-^ and H_2_O_2_, resulted in shorter primary roots compared to their counterparts without H_2_O_2_, PAW1 and 4, respectively (Supplemental Fig. 1A). Therefore, PAW1, 4, and 5 showed increased total root growth compared to NO_3_^-^ and NO_3_^-^ plus H_2_O_2_ despite PAW5 having a reduced primary root growth (Fig. 1B). To determine whether differences in total root length were due to secondary root initiation or elongation, we measured lateral root density in PAW-treated plants. PAW-treated plants showed similar or reduced lateral root density compared to both the NO_3_^-^ and NO_3_^-^ plus H_2_O_2_ controls. Given that seedlings treated with PAW1, PAW4, or PAW5 have longer roots without significant increases in lateral root density, increases in root lengths appear to be due to elongation and not increased lateral root initiation (Supplemental Fig. 1B). PAW4 and PAW5 were chosen as the optimal PAW chemistries for further experiments due to the increased elongation seen in treated seedlings.

**Figure 1.**
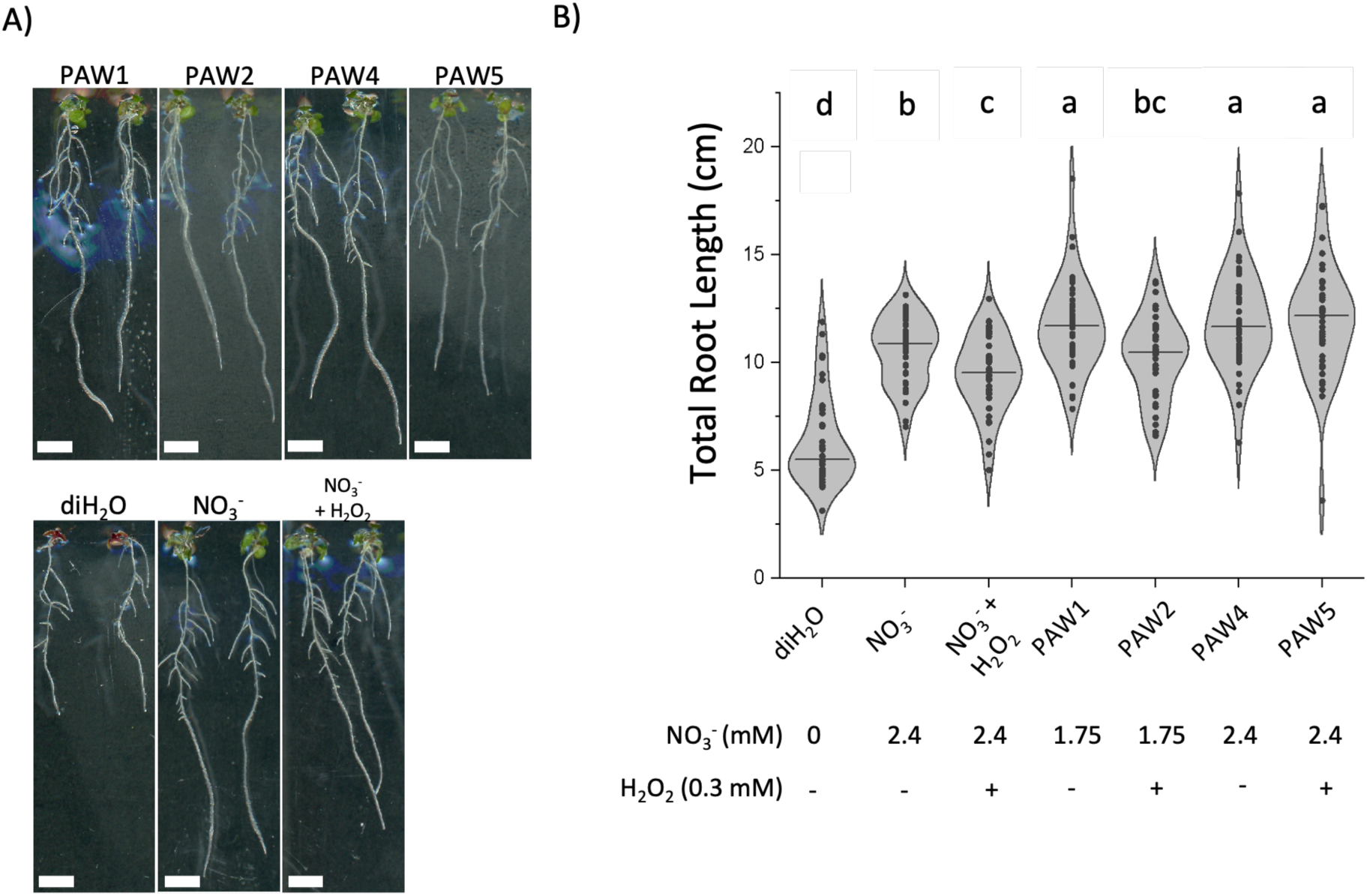
Optimal PAW treatment promotes total root elongation of Arabidopsis seedlings. A) Three-day old Arabidopsis seedlings were transferred to N-free media supplemented with PAW1, PAW2, PAW4 or PAW5, diH_2_O, 2.4mM NO_3_^-^ or 2.4mM NO_3_^-^ and 0.15mM H_2_O_2_. Seedlings were imaged after 5 days. B) The root length of primary and secondary roots combined was measured from seedlings treated as in A). A one-way ANOVA with Tukey’s multiple comparison test was performed. N= 40 seedlings.

**Table 1.** Concentrations of NO_3_^-^ and H_2_O_2_ in treatment solutions. Expected concentration of NO_3_^-^ and H_2_O_2_ from each PAW or control solution after incorporation into gel media. The solutions consist of the Low N, NO_3_^-^, and H_2_O_2_ + NO_3_^-^ treatments were mixed from NO_3_^-^ salts and diluted H_2_O_2_ to match the concentrations of the PAW treatments.

| ID | NO<sub>3</sub><sup>-</sup> (mM) | H<sub>2</sub>O<sub>2</sub> (mM) |
| --- | --- | --- |
| diH<sub>2</sub>O | 0 | 0 |
| Low N | 0.25 | 0 |
| NO<sub>3</sub><sup>-</sup> | 2.4 | 0 |
| H<sub>2</sub>O<sub>2</sub> + NO<sub>3</sub><sup>-</sup> | 2.4 | 0.15 |
| PAW1 | 1.75 | 0 |
| PAW2 | 1.75 | 0.15 |
| PAW4 | 2.4 | 0 |
| PAW5 | 2.4 | 0.15 |

### ROS Present in PAW Inhibit Root Growth in Arabidopsis Seedlings

Previous research suggested that RONS in PAW solutions enhance plant growth (Rashid et al., 2021; Rathore et al., 2022), but both PAW2 and PAW5 showed shorter primary roots compared to their counterparts without H_2_O_2_. Moreover, the NO_3_^-^ content in PAW alone does not explain the differing response of PAW-treated plants as PAWs with different levels of H_2_O_2_ showed different levels of root growth. RONS are still a major component of PAW (Gierczik et al., 2020; Iseni et al., 2016; Kučerová et al., 2019; Traylor et al., 2011) with H_2_O_2_ being the most concentrated and stable ROS present in the PAW2 and PAW5. To further investigate the role of H_2_O_2_ on the plant response to PAW, we measured root growth in *Arabidopsis* seedlings (Col-0) treated with 1.75 mM NO_3_^-^ and varying concentrations of H_2_O_2_. Seedlings treated with up to 0.25 mM H_2_O_2_ appeared healthy with limited visual indications of stress (Fig. 2A), but these seedlings presented 18% shorter primary roots on average compared to seedlings treated with the NO_3_^-^ plus 0.0 mM H_2_O_2_ control. Even treatments as low as 0.05 mM H_2_O_2_ resulted in a 14% reduction in primary root length when compared to the NO_3_^-^ without H_2_O_2_ (Fig. 2B). Ultimately, H_2_O_2_ in NO_3_^-^containing solutions resulted in inhibition of primary root growth.

**Figure 2.**
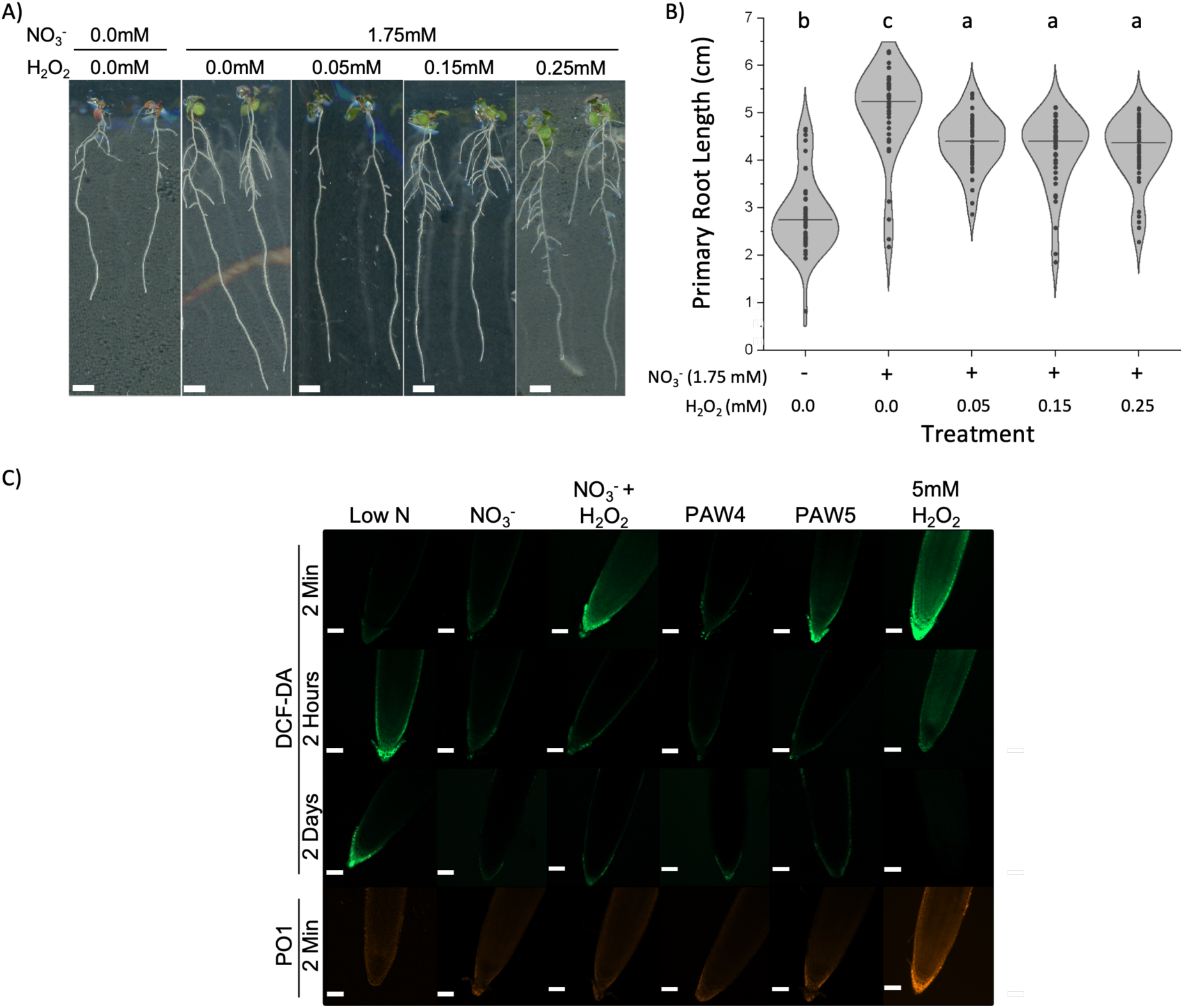
H_2_O_2_ was detrimental to root length. A) Three-day old Arabidopsis seedlings were transferred to N-free media supplemented with 0 (diH_2_O) or 2.4mM NO_3_^-^ and 0-0.25 mM H_2_O_2_ and incubated for 5 days. B) The primary root length was measured from seedlings treated as in A). Data shows results of one-way ANOVA with Tukey’s multiple comparison test. N= 40 seedlings. C) Three- day old seedlings were incubated in N-free media supplemented with 0.25 mM NO_3_^-^ (Low N), 2.4 mM NO_3_^-^ (NO_3_^-^) with or without 0.15 mM H_2_O_2_, PAW4, PAW5, or 2.4 mM NO_3_^-^ with 5 mM H_2_O_2_ for 2 minutes, 2 hours, or 2 days and then stained with 2′-7′ dichlorodihydrofluorescein diacetate (DCF-DA) or Peroxy Orange 1 (PO1). Root tips were imaged by confocal microscopy (scale = 100 µm).

The H_2_O_2_ in PAW5 inhibited primary root elongation but other ROS potentially present in PAW could contribute to the elongated root response to PAW4. Cells in the root tip and surrounding tissue are responsible for controlling root elongation by controlling rates of cell division and cell elongation (Bouguyon et al., 2012; Maghiaoui et al., 2020). We therefore tested whether short PAW treatments elicited the accumulation of different ROS in root tips using the fluorescent dyes 2′-7′ dichlorodihydrofluorescein diacetate (H_2_DCFDA), a general intracellular ROS probe (Kuběnová et al., 2023; Martin et al., 2022), and Peroxy Orange 1 (PO1), a specific probe for intracellular H_2_O_2_ (Dickinson et al., 2010; Martin et al., 2022). Seedlings transferred to N deficient media resulted in ROS accumulation in Arabidopsis root tips after 2 hours as detected by H_2_DCFDA staining (Fig. 2C), as seen in other studies (Shin et al., 2005; Zhu et al., 2016). No apparent changes in H_2_DCFDA fluorescence were detected up to 2 days after transfer of seedlings to N-replete media (NO_3_^-^ control) (Fig. 2C). In contrast, seedlings transferred to media containing NO_3_^-^ plus H_2_O_2_ showed increased H_2_DCFDA fluorescence along the root cap and root tip epidermis but only at the 2 minutes time point (Fig. 2C). At later time points, reduced fluorescence was found across all treatments except the Low Nitrogen (Low N) control. The activity of scavenging enzymes including catalases (Bai et al., 1999; Bi et al., 2009) may explain the reduction in H_2_DCFDA fluorescence at the later times points. We found that roots treated with PAW4-containing media exhibit similar pattern and intensity of H_2_DCFDA staining compared to the NO_3_^-^ control at all time points (Fig. 2C), suggesting the absence of other ROS components unaccounted for in PAW4. On the other hand, PAW5-treated seedlings showed similar staining as the NO_3_^-^ + H_2_O_2_ control with stronger H_2_DCFDA after 2-minutes of seedling transfer but reduced fluorescence in later timepoints. This increase in H_2_DCFDA fluorescence 2 minutes after treatment with PAW5 or the NO_3_^-^ plus H_2_O_2_ control may be attributed to endogenous ROS production in response to the elevated H_2_O_2_ in the media, similar to a stress response (Guler & Pehlivan, 2016; Kuběnová et al., 2023; Savvides et al., 2016; Tognetti et al., 2017). A high (5 mM) H_2_O_2_ + NO_3_^-^ treatment was used to determine successful incubation with the probe and resulted in high levels of DCF fluorescence throughout multiple cell layers of the root after 2 minutes. H_2_DCFDA staining also decreased in these seedlings after 2 hours or 2 days of treatment. No differences in Peroxy Orange 1 (PO1) fluorescence intensity were detected in seedlings treated with PAW4, PAW5 or the corresponding NO_3_^-^ controls with or without H_2_O_2_. PO1 staining was only detected in seedlings exposed to the 5 mM H_2_O_2_ control. This result suggests that the accumulation of endogenous H_2_O_2_ is not the primary ROS produced under PAW5 and the NO_3_^-^ plus H_2_O_2_ treatments. It is important to note that as a boronate-based probe, PO1 can react slower than a probe like H_2_DCFDA which can cause short-term ROS accumulation to appear weaker as the probe needs more time to respond (Martin et al., 2022; Winterbourn, 2014).

### PAW5 pre-treatment confers some protection to plants under heat response

H_2_O_2_ has been linked to priming in plants, which can confer resistance to abiotic stressors (Guler & Pehlivan, 2016; Rodríguez-Ruiz et al., 2019; Savvides et al., 2016; Zhu et al., 2019). The different concentrations of ROS in PAW solutions allowed us to determine if PAW could provide a priming effect to protect against elevated temperatures in Arabidopsis. Three-day old seedlings (Col-0) were transferred to PAW or NO_3_^-^ control media and incubated at 22°C for 1 day before moving them to 22°C or 30°C for an additional 4 days. The 22°C control set of seedlings reflected root growth seen in prior experiments where PAW5 treated seedlings showed similar or reduced primary root length compared to both PAW4 and the NO_3_^-^ control (Fig. 3A, B). Seedlings treated with high or low NO_3_^-^ showed similar root lengths at 30°C when compared within this temperature group (Fig. 3B), suggesting that N availability did not impact the response to elevated temperatures. PAW4-treated seedlings showed similar root growth compared to NO_3_^-^ controls, indicating that PAW4 does not protect plants against heat stress (Fig. 3B). Plants undergoing PAW5 treatment, which contains 0.15 mM of H_2_O_2_, showed a significant increase in primary root length compared to the NO_3_^-^ plus H_2_O_2_ control at the end of the elevated temperature period, with a 46% increase in primary root length (Fig. 3B). PAW5-treated seedlings had 27% longer primary roots compared to PAW4-treated seedlings after 4 days of heat stress (Fig. 3B), which is opposite to the effects of these PAW treatments at 22°C (Supplemental Fig. 1A). This suggests that ROS in PAW5 elicit a improved response to elevated temperatures compared to other treatments in that group. However, this cannot be fully explained by the H_2_O_2_ content in PAW5, as the NO_3_^-^ plus H_2_O_2_ control were not sufficient to cause the same improved growth under elevated temperatures.

**Figure 3.**
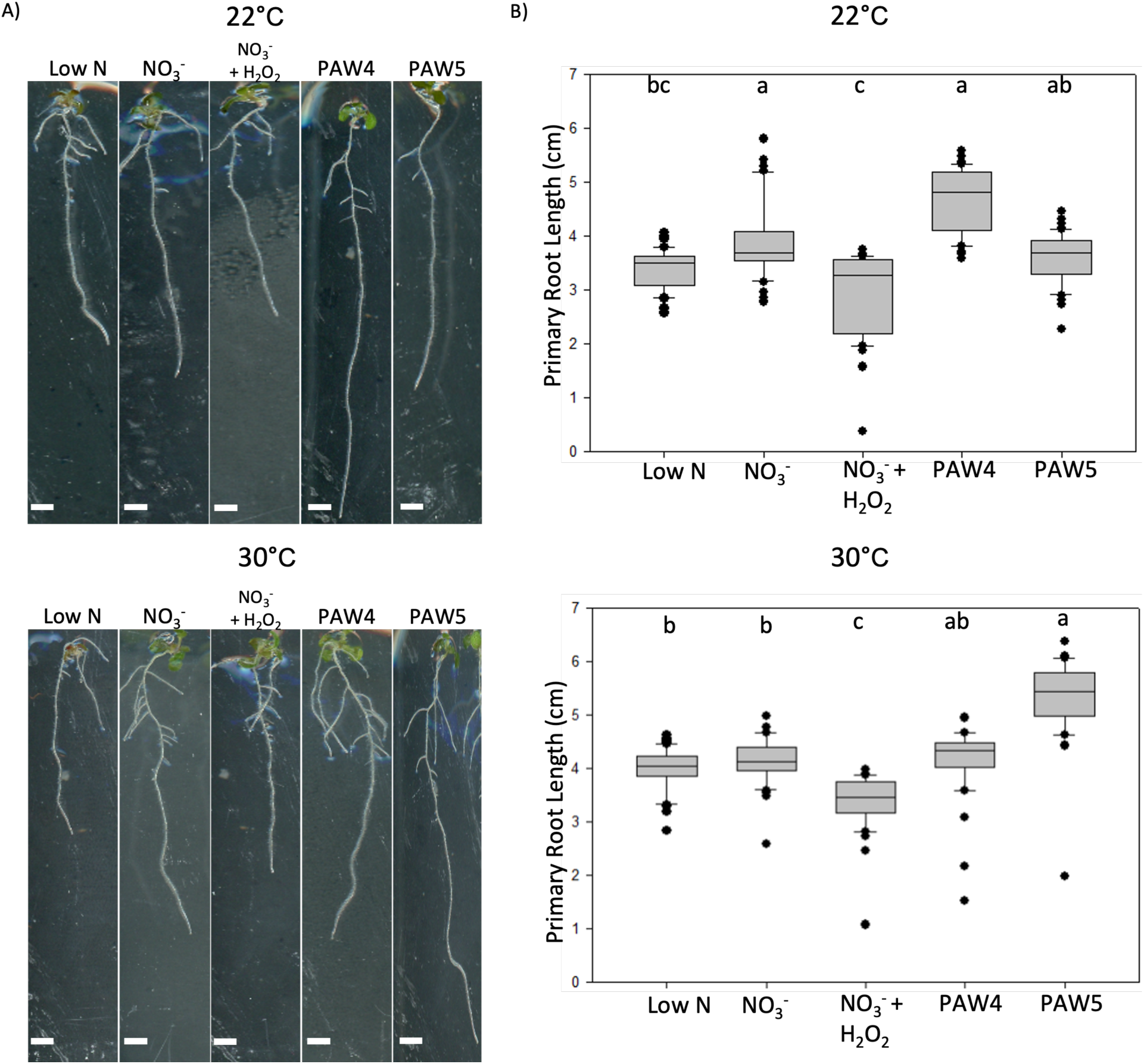
PAW5 Confers Resistance to Heat Stress. A) Three-day old Arabidopsis seedlings were transferred to N-free media 0.25 mM NO_3_^-^ (Low N), 2.4 mM NO_3_^-^ (NO_3_^-^) with or without 0.15 mM H_2_O_2_, PAW4 or PAW5 and incubated for 1 day at 22°C. Plates were either kept at 22°C (top) or incubated at 30°C (bottom) for 5 days. B) Primary root length was measured from seedlings at the end of the 22°C (top) or 30°C (bottom) incubation as described in A). A one-way ANOVA with Tukey’s multiple comparison test was performed. N= 40 seedlings.

### PAW treatment in seedlings shows no detectable change in hormone accumulation in root tissue

Seedlings exhibit increased root elongation under treatment with PAW4; however, the underlying cause of this elongation is not yet known. We investigated whether PAW treatment resulted in changes in plant hormone signaling in root tissue by visualization of fluorescent markers. Markers for auxin, ethylene, and cytokinin response were selected due to the role of these hormones in controlling root morphology under varying NO_3_^-^ levels (Bouguyon et al., 2015; Bouguyon et al., 2012; Krouk et al., 2010). The DR5::GFP marker is a transcriptional fusion of the synthetic DR5 promoter with GFP such that GFP accumulates in cells where auxin signaling is active (Ulmasov et al., 1997). A EBS::YPet marker was used to visualize ethylene response as the EBS promoter functions downstream of ethylene perception and signaling (Fernandez-Moreno & Stepanova, 2020). The TCSn::GFP marker was used to compare the response to cytokinin, as the TCSn promoter is responsive to that hormone (Liu & Müller, 2017; Zürcher et al., 2013). Arabidopsis seedlings expressing each marker were treated with PAW4, PAW5, or the NO_3_^-^ control for two days and seedlings were imaged by confocal microscopy (Fig. 4). All treatments with the DR5::GFP resulted in similar pattern of GFP fluorescence in the root meristematic region and in the innermost cell files of the root cap (Ulmasov et al., 1997; Vicente-Agullo et al., 2004). Similarly, all treated EBS::YPet roots showed similar patterns of YPet fluorescence concentrated at the root tip and the epidermis as previously reported (Herrera-Rodríguez et al., 2022; Stepanova et al., 2007). Finally, no differences between treatments were detected with the TCSn::GFP lines with fluorescence signal detected in the root epidermis approaching the apex of the root (Liu & Müller, 2017; Steiner et al., 2020). These results suggest that PAW4 and PAW5 treatments do not induce changes in auxin, ethylene, or cytokinin signaling pathways in the root compared to the NO_3_^-^ control.

**Figure 4.**
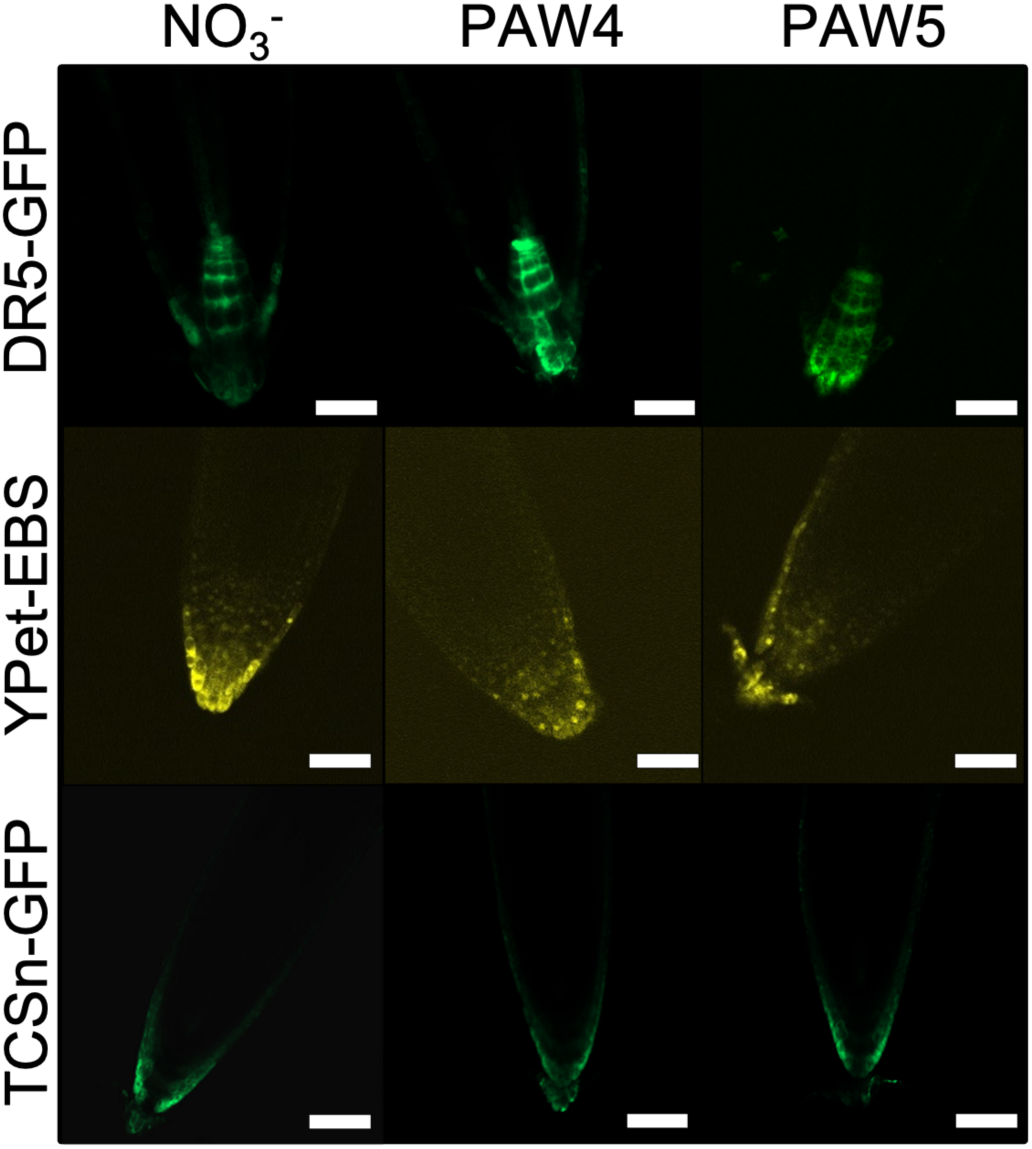
PAW-treatment does not alter the pattern of hormone response markers in roots. *Arabidopsis* seedlings expressing either the DR5::GFP, EBS::Ypet, or TCSn::GFP biomarkers were grown for 3 days on full nutrient media and then transferred to N-free media supplemented with 2.4mM NO_3_^-^ (control), PAW4 or PAW5. Root tips were imaged by confocal microscopy after 2 days of treatment. Scale = 100 µm.

### PAW treatment induced Nitrogen-responsive gene expression

Small increases in total root length were found for seedlings treated with PAW4 when compared to an equivalent NO_3_^-^ control. This led to the hypothesis that PAW treatment may result in changes in gene expression when compared to the N control. Wild-type three-day old seedlings were treated with PAW4 or an equivalent amount of NO_3_^-^ for 8 days as described in previous experiments. RNA was extracted separately from shoots and roots from at least 20 seedlings per sample and sequenced using Next-Generation Sequencing. Sequence data was used to compare gene expression between PAW-treated tissues with the NO_3_^-^ controls. We should note that Principal Coordinate Analysis (PCA) of root samples indicated that one root sample each from PAW and control treatments did not cluster with the like samples (Supplemental Figure 2). The analysis described below included these samples due to the limited number of replicates.

Differential expression analysis between PAW-treated tissues compared to the NO_3_^-^ control samples included genes with FDR p-values less than or equal to 0.01 and at least a 2-fold change or greater at a log_2_ transformation. A total of 352 genes were upregulated, while 243 downregulated in PAW-treated roots compared to the controls (Fig. 5A). A gene ontology (GO) analysis was carried out to identify biological pathways that may be altered by PAW exposure. Genes associated with nitrate assimilation and related metabolic pathways showed significant enrichment in PAW-treated root tissue when compared to the control (Fig. 5B). Additionally, genes associated with responses to oxygen-containing compounds such as ROS showed approximately 50 genes with moderate fold enrichment (Fig. 5B). In the downregulated genes in roots, genes associated with H_2_O_2_ metabolism and catabolism exhibited significantly lower fold enrichment compared to the NO_3_^-^ control as well as showing high -log_10_(FDR) values (Fig. 5B). Notably, genes associated with glutamine catabolic processes showed a low proportion of the downregulated genes, however, still exhibited significance (Fig. 5B). Glutamine is highly regulated in the plant and control can also be varied by NH_4_^+^ availability and perception. The catabolism of glutamine can result in glutamate, which can be recycled to form other N compounds (Miflin & Habash, 2002).

**Figure 5.**
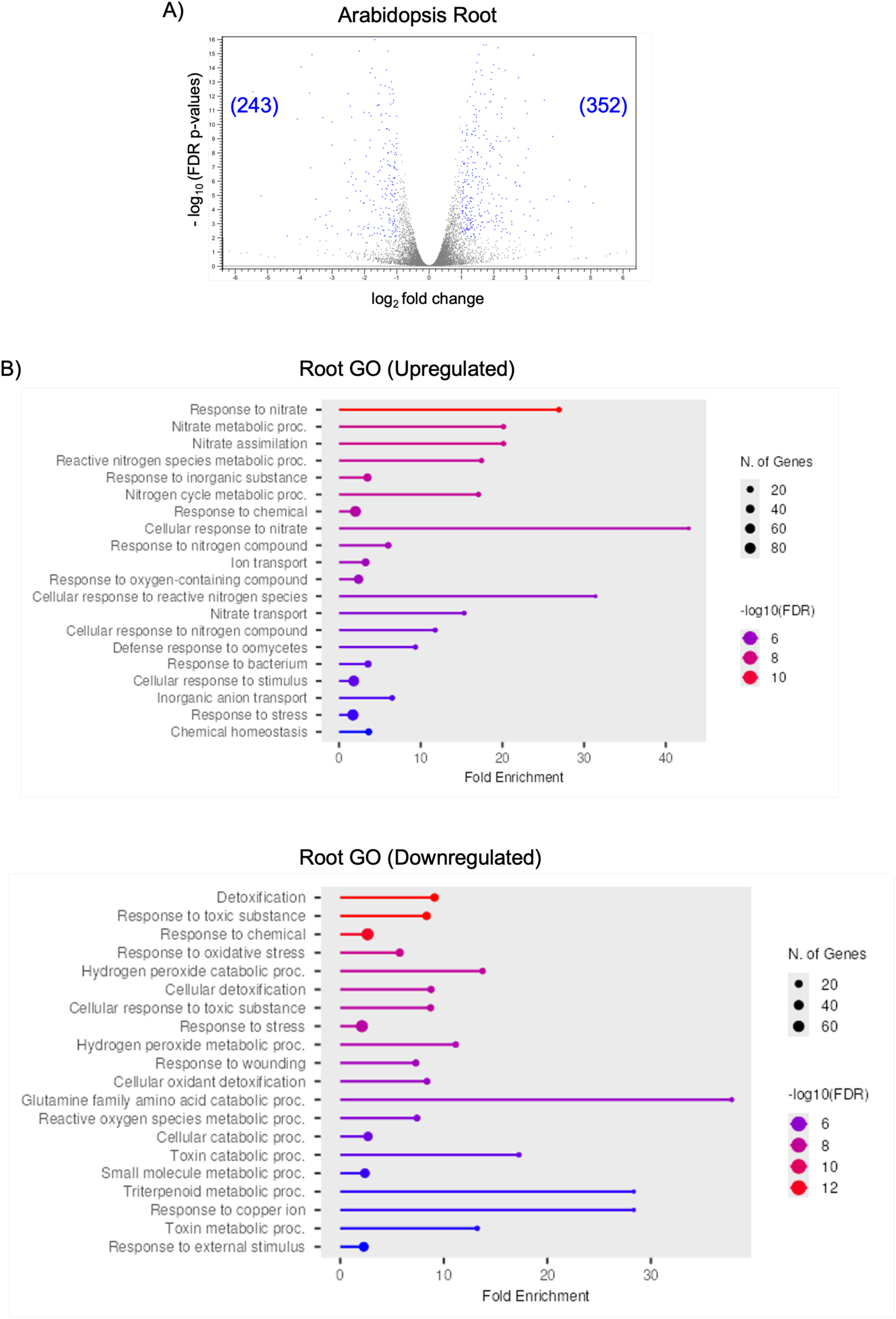
PAW treatment results in differential expression in genes associated in responses to N and ROS in Arabidopsis roots. Three-day old Arabidopsis seedlings were transferred to N-free media supplemented with PAW4 or 2.4mM NO ^-^ as a control. Roots were harvested one week later and RNA was extracted and sequenced. A) Volcano plots of differentially-expressed genes in roots between PAW-treated and NO ^-^-treated controls. Numbers represent the number of differentially expressed genes (blue dots) with p-values less than or equal to 0.01 and 2-fold or greater absolute fold change. B) Gene Ontology (GO) analysis characterized by biological processes of differentially expressed genes that were up- or down-regulated in the root.

In the shoot samples, 309 genes were upregulated and 492 were downregulated in the PAW samples compared to the controls (Fig. 6A). Upregulated genes were enriched in pathways associated with regulation of flavonoid biosynthesis and responses to nitrogen in shoot tissue (Fig. 6B). Flavonoids serve a variety of roles in the plant including scavenging of ROS in plant cells (Di Ferdinando et al., 2012). This data suggests that seedlings treated with PAW exhibited an altered response to nitrate which can affect uptake and rate of assimilation. Additionally, while PAW4 was intended to have limited concentrations of ROS, the analysis suggests that plants treated with PAW may be experiencing a response to oxidative compounds. In the downregulated genes, genes associated with low O_2_ levels and stress responses were found to be the most significantly impacted (Fig. 6B).

**Figure 6.**
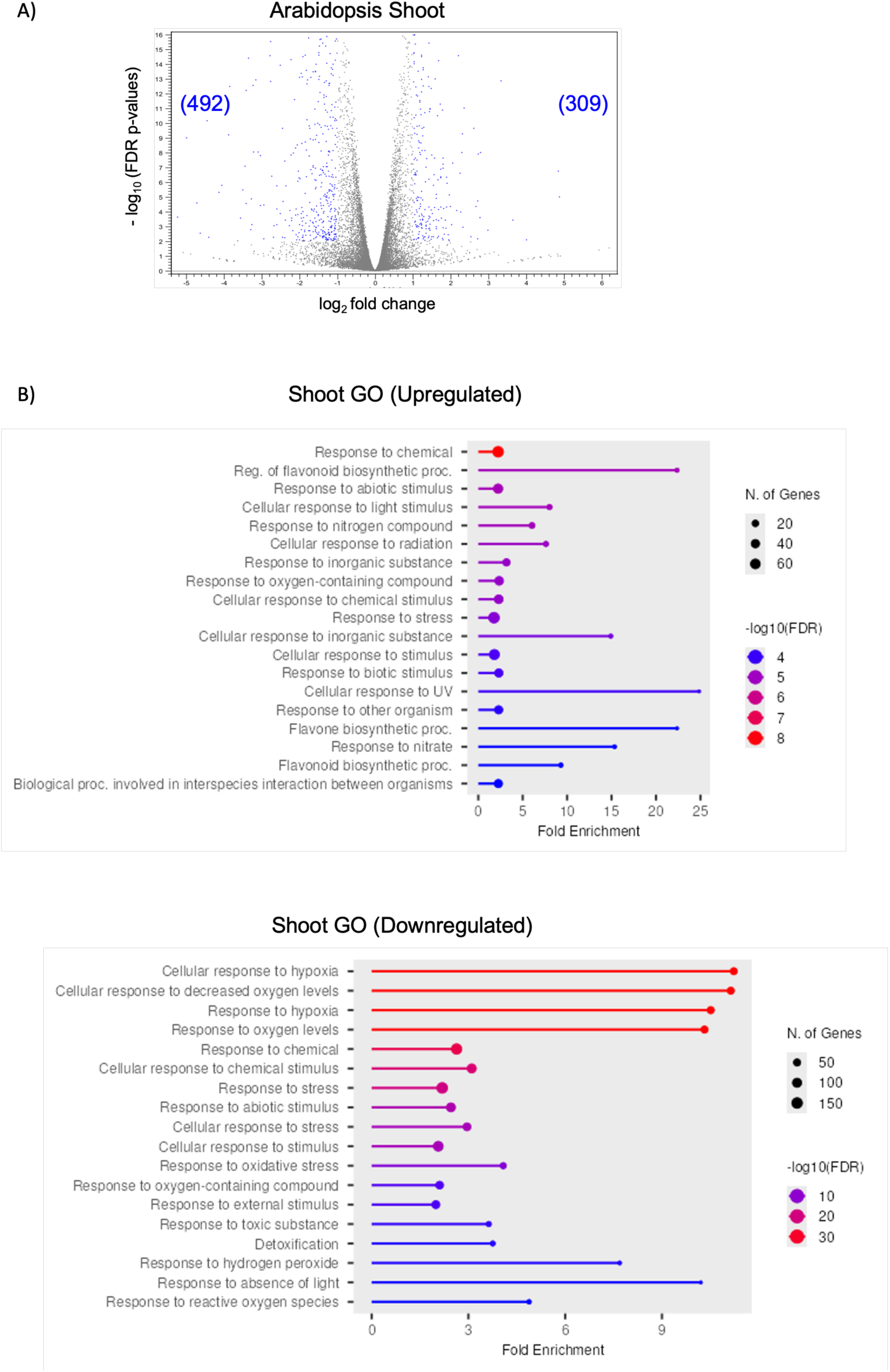
PAW treatment results in differential expression in genes associated in responses to N and ROS in Arabidopsis shoots. Three-day old Arabidopsis seedlings were transferred to N-free media supplemented with PAW4 or 2.4mM NO_3_^-^ as a control. Shoots were harvested one week later and RNA was extracted and sequenced. A) Volcano plots of differentially-expressed genes in shoots between PAW-treated and NO_3_^-^-treated controls. Numbers represent the number of differentially expressed genes (blue dots) with p-values less than or equal to 0.01 and 2-fold or greater absolute fold change. B) Gene Ontology (GO) analysis characterized by biological processes of differentially expressed genes that were up- or down-regulated in the shoot.

### PAW4 is an effective fertilizer for soil-grown plants

While PAW treatment induces changes in root morphology in Arabidopsis 5-day old seedlings (Ka et al., 2021), little is known about the effects of prolonged treatment with PAW. We next evaluated the growth of Arabidopsis plants treated for up to 5 weeks with concentrated PAW4 (4.8 mM NO_3_^-^) because of its optimal performance compared to PAW5. Plants were regularly watered with diH_2_O and biweekly with a ¼ x Hoagland solution without N to provide all other essential nutrients. Plants were watered weekly with one of the treatment solutions, the NO_3_^-^ control or concentrated PAW4, as the only available source of N. After a 5-week growing period, plants were harvested to measure fresh and dry biomass of both shoot and root.

All plants developed healthy rosettes and inflorescences after 5 weeks (Fig. 7A). No differences were detected in rosette area between PAW4-treated plants and those of the NO_3_^-^ control (Fig. 7B), but PAW4-treated plants showed a reduction in shoot fresh mass (Fig. 7C). PAW4-treated plants also showed reduced shoot dry biomass when compared to plants treated with the NO_3_^-^ control (Fig. 7D). Interestingly, the PAW4-treated plants showed comparable dry root biomass to plants treated with the NO_3_^-^ control (Fig. 7E). However, taking the total dry biomass by combining both dry aboveground and root biomass yielded statistically similar biomass between the two treatments (Fig. 7F). Equivalent growth between the two treatments was also documented by their similar root:shoot ratio (Fig. 7G). Despite reduced shoot dry biomass and similar root dry biomass of PAW4-treated plants, the root:shoot ratio suggests no difference of resource allocation between plants in either treatment. These results combined suggest that PAW4-treatment results in comparable growth and overall dry biomass as plants treated with the NO_3_^-^ control and support the use of PAW as a potential alternative to existing NO_3_^-^ fertilizers.

**Figure 7.**
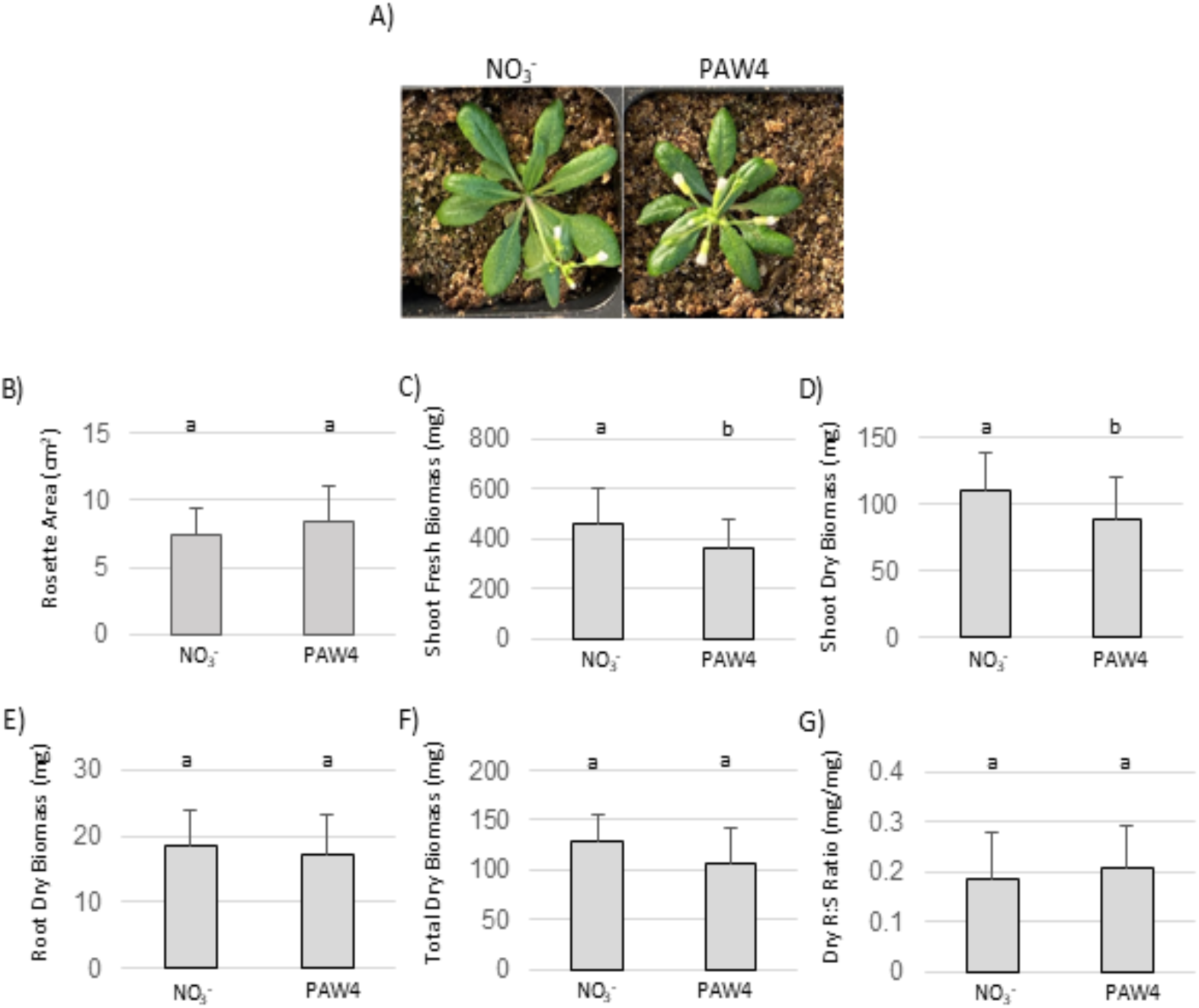
PAW is a good substitute for NO_3_^-^ fertilizer. A) Three-day old wild-type Arabidopsis seedlings were grown were transplanted to soil without N and fertilized with either 4.8mM NO_3_^-^ or PAW4 (4.8mM NO_3_^-^) for 5 weeks. One representative image is shown. B) Rosette area was measured from top images with ImageJ. C-G) Plants were harvested after 5 weeks of treatment and shoots and roots were used to measure shoot fresh biomass (C), shoot dry biomass (D), root dry biomass (E), Total dry biomass (F) and root:shoot ratio G). N = 20 plants per treatment. Significance denotes p < 0.05 in Student’s T-test.

## DISCUSSION

### PAW Can be Used as an Alternative for N Fertilizer

Nitrogen is a macronutrient for plants because it is a major component of nucleic acids and proteins. Plants primarily take up N through their roots, sourcing the inorganic forms NO_3_^-^ and NH_4_^+^ from the soil environment (Hachiya & Sakakibara, 2017). Synthetic fertilizers containing NO_3_^-^ and NH_4_^+^ have become globally the most abundant type of N fertilizer over the last few decades (Lassaletta et al., 2014). Methods to produce these fertilizers are unsustainable, and they have an additional carbon footprint through transport and the supply chain (M. Wang et al., 2021). There is an urgent need for the development of sustainable fertilizers that can meet growing agricultural demands, which PAW from non-thermal plasmas has the potential to meet (Chen et al., 2018). However, diverse technologies for non-thermal plasma generation and water activation are still under development, and therefore, a unified standard in PAW production is not yet established (Ranieri et al., 2021; Robinson & Stapelmann, 2024).

PAW has been proposed as an effective source of N for plant growth (Kučerová et al., 2019; Li et al., 2021; Ndiffo Yemeli et al., 2021; Rashid et al., 2021; Rathore et al., 2022; Škarpa et al., 2020). The most important contribution of PAW comes from its NO_3_^-^ content as substituting NO_3_^-^ solutions for PAW completely restores root growth (as compared to the water controls without N). PAW treatments with identical NO_3_^-^ content resulted in longer roots when compared to the NO_3_^-^ controls at the seedling stage indicating the presence of additional molecules with growth-promoting activity. These differences are certainly mild and unlikely due to H_2_O_2_ because this ROS was mostly inhibitory in controlled growth experiments in seedlings. Our results are consistent with previous reports that showed enhanced germination and longer roots in PAW-treated Arabidopsis seedlings (Bafoil et al., 2019; Cui et al., 2019; Ka et al., 2021).

The transcriptome analysis showed that PAW treatment induced the expression of genes associated with N uptake and assimilation in roots. Genes from the NRT2 family such as NRT2.1, NRT2.6, and NRT2.4 were amongst the significantly upregulated genes. NRT2 family transporter proteins are characterized as high-affinity NO_3_^-^ transporters, often preferentially expressed in the root tissue, and function in uptake from the soil environment (Okamoto et al., 2003; Orsel et al., 2002). NRT2.1 is known as a nitrate-inducible transporter that is highly expressed in the root when Arabidopsis is exposed to NO_3_^-^ (Okamoto et al., 2003). The upregulation of these genes suggests that seedlings treated with PAW are primed to more readily transport NO_3_^-^ from the soil environment. Furthermore, genes associated with NO_3_^-^ metabolism and NO_3_^-^ assimilation were upregulated in these roots, which suggests increased perception and response to internal NO_3_^-^ levels in root cells (Lam et al., 1996; Stitt, 1999; Wang et al., 2000). Overall, the increased expression of these genes may contribute to the mild root length increases noted at the seedling stage. While this differential expression exhibited by plants treated with PAW may suggest PAW treatment improves NO_3_^-^ uptake and assimilation, these responses may only be short term. PAW4 treatment of mature plants in soil resulted in similar root growth as the NO_3_^-^ controls. This suggests that the enhanced root elongation effect of PAW detected in seedlings may be restricted to early developmental stages or to highly controlled environments such as the sterile Arabidopsis growth media. Treatment of plants with PAW4 for 5 weeks in soil resulted in plants with similar morphology and development compared to those watered with NO_3_^-^ control solution. Overall, there were no significant effects of PAW treatment in terms of total dry biomass and rosette area when compared to the NO_3_^-^ controls, indicating that plants respond to PAW in a similar manner as regular fertilizer in long-term experiments. Our results showed PAW to be a sufficient fertilizer and alternative to traditional NO_3_^-^ fertilizers for Arabidopsis seedlings and 5 week old rosettes. Further studies focused on the upregulation of genes associated with NO_3_^-^ uptake are required to determine if treatment with trace ROS like those present in PAW4 may have benefits beyond nitrogen fertilization.

Many recent reports highlight dramatic differences in PAW-treated plants when compared to plants treated with tap water or deionized water alone. In barley for example, certain PAW formulations resulted in up to 37% increase in fresh biomass compared to the water control over a 4-week period (Ndiffo Yemeli et al., 2021), while other PAWs have increased yield of rice by 16.7% after a full growing season (Rashid et al., 2021). Maize showed a 13.1% dry biomass increase; however, that was through foliar sprays (Škarpa et al., 2020). Pea plants also showed improved root and shoot growth with increases up to 38% and 95% respectively, compared to a tap water control (Rathore et al., 2022). Most of these effects are likely attributed to the nitrogen content in PAW compared to the absence of nitrogen in the control treatment. However, these studies do not directly evaluate the potential of PAW as an alternative to traditional nitrogen fertilizers. The lack of an equivalent NO_3_^-^ control, as well as the variability in chemical composition of PAWs from different plasma devices, make comparisons between different studies challenging. Moreover, very few studies specify the soil composition or tightly control for N content in soil. Most commercial soil or substrate formulations are likely amended with sufficient fertilizer for minimal plant growth, even if not specified in the label (Dalling et al., 2013; Williams, 1995), and this can seriously affect interpretation of PAW effects on plant growth. Our study used unamended commercial peatmoss, vermiculite, and perlite as raw materials to make soil substrates where NO_3_^-^ or PAW solutions were the only sources of plant-available N. Unlike previous PAW studies where plants in soil media were watered with PAW (Kučerová et al., 2019; Lamichhane et al., 2021; Rathore et al., 2022), plants did not survive beyond 3 weeks when watered with diH_2_O plus Hoagland minus N alone. We propose that future PAW studies include NO_3_^-^ equivalent solutions to allow for comparisons between studies and to highlight the consistency of PAW as an alternative fertilizer.

### ROS in PAW Inhibits Growth Under Ideal Growth Conditions

Previous reports have proposed that ROS in PAW contribute to growth enhancement beyond that of nitrate (Rashid et al., 2021; Rathore et al., 2022). By generating PAW solutions that differed on the concentration of NO_3_^-^ and the presence of H_2_O_2_, we were able to study the independent contributions of N and ROS. Exogenous ROS can be detrimental to plant health, especially at high concentrations, whereas low levels of ROS were proposed to enhance plant growth (Beckman & Ames, 1997; Berlett & Stadtman, 1997; Berrios & Rentsch, 2022; Singh et al., 2016). The primary root lengths of seedlings treated with both PAW types (1.75 and 2.4 mM NO_3_^-^) were shorter when H_2_O_2_ was present, indicating that even low concentrations of this ROS inhibited root growth. This was corroborated with controlled experiments with media containing as low as 0.05 mM of H_2_O_2_ which resulted in shorter roots in seedlings exposed to the same levels of N. These results suggest that PAW chemistries containing the lowest levels of ROS might be a better alternative for fertilizer substitution. Similar results previously reported that increased concentration of H_2_O_2_ in PAW result in oxidative stress, inhibition of seedling growth and shorter primary roots (Cui et al., 2022; Cui et al., 2019; Ka et al., 2021; Panngom et al., 2018; Zhou et al., 2018). Arabidopsis specifically showed a reduction in root elongation as concentrations of H_2_O_2_ increased, resulting in a 28% reduction in total root length after 7 days (Ka et al., 2021). These results together with those reported in this study demonstrate that H_2_O_2_ does not provide any added benefit in PAW under optimal growing conditions.

An array of different ROS may accumulate within PAW (Iseni et al., 2016), but are difficult to measure (Ran et al., 2024). Theoretically, reactive nitrogen species (RNS) like peroxynitrite (ONOO^-^) or other ROS like superoxide (O_2_ ^−^) may be present in the PAW (Iseni et al., 2016; Kučerová et al., 2019; Tachibana & Nakamura, 2019), but at the current time the presence and concentrations of these species is unknown due to limitations in methods to measure these species cost-effectively. We used fluorescent probes (Martin et al., 2022) as proxies to elucidate the effect of PAW4 and PAW5 on the plant root redox state. H_2_DCFDA responds generally to oxidation from endogenous ROS, which works effectively in characterizing ROS response in *Arabidopsis* (Kuběnová et al., 2023; Martin et al., 2022). Both PAW4 and PAW5 treatments resulted in similar patterns of H_2_DCFDA staining when compared to their corresponding NO_3_^-^ or NO_3_^-^ plus H_2_O_2_ controls. We interpret this result as evidence that the main ROS in PAW is likely H_2_O_2_, and that no ROS are present in PAW4 at concentrations that would elicit a plant response. This is further reinforced with PO1 staining, which specifically reports the accumulation of endogenous H_2_O_2_. From the 2-minute time point, both PAW 4 & 5 treatments and their corresponding controls containing NO_3_^-^ with or without H_2_O_2_ showed no difference in their response. The only response can be found from the root treated with a high (5 mM) concentration of H_2_O_2_. It is important to note that the NO_3_^-^ plus low (0.15 mM) concentration of H_2_O_2_ control showed no PO1 staining where the only difference between the two controls is the concentration of H_2_O_2_. This can be interpreted by a difference in sensitivity between H_2_DCFDA and PO1, but studies where both H_2_DCFDA and PO1 were used did not report major differences in sensitivity between these dyes (Gayomba & Muday, 2020; Watkins et al., 2017). PO1 is a boronate-based probe, which means it can respond more specifically to H_2_O_2_ presence than H_2_DCFA. However, this type of probe can react slower than H_2_DCFDA (Martin et al., 2022; Winterbourn, 2014). If the peak of the intracellular H_2_O_2_ produced due to PAW treatment has a limited window of time on the scale of minutes, a probe like PO1 may be too slow and report a weaker signal. It also suggests that the response to 0.15 mM H_2_O_2_ elicits a short-term response of endogenous ROS production where H_2_O_2_ is produced in low concentrations or is swiftly scavenged by antioxidants like catalase. This may explain why PAW treatment did not yield a strong response to ROS and PAW under these experimental conditions like the response found with H_2_DCFDA.

### PAW May Confer Stress Resistance Through Priming with ROS

PAW containing ROS may potentially provide stress protection to crop plants in the form of “priming” that lessens the impact of stressors (Mittler & Blumwald, 2015; Rodríguez-Ruiz et al., 2019; Savvides et al., 2016). Some work has found that direct treatment of seeds with non-thermal plasma conferred benefits to seedlings undergoing osmotic and saline stress (Bafoil et al., 2019). PAW treatment also enhanced cold-tolerance in tomato (Li et al., 2021) and biotic stress resistance in grape vines (Zambon et al., 2020). It has been theorized that the RONS present in PAW are substantial enough to initiate the priming effect. We focused on high temperature stress, which is becoming a more relevant stress for agriculture as global climate patterns continue changing at an accelerated rate (Ainsworth & Ort, 2010). A PAW5 pre-treatment seemed to protect Arabidopsis seedlings during a short-term high temperature stress, but PAW4 did not. This may be due to the presence of RONS in PAW5 (Fedina et al., 2009; Uchida et al., 2002). It is important to note that a similar concentration of H_2_O_2_ was insufficient to provide the same benefit as PAW5, suggesting the presence of other molecular species in PAW5 that may enhance the priming effect. Overall, these results highlight the potential benefit of generating PAW with different levels of ROS to facilitate plant growth under stress and non-stress conditions.

## ACKNOWLEDGMENTS

This work was supported by the NCSU Game-Changing Research Incentive Program for Plant Science Initiative (GRIPS4PSI to K.S.), the Research Capacity Fund (HATCH), project award no. 7005574, from the U.S. Department of Agriculture’s National Institute of Food and Agriculture, US Department of Education GAANN fellowship to J.K. and an NC Space Grant award to J.K. We thank Katherine Danz for technical assistance, Anthony Postiglione and Brian Whipker for advice and Anna Stepanova for materials. The authors also acknowledge the Rojas-Pierce laboratory for valuable discussions

## Conflict of Interest

The authors of this work declare that there is no conflict of interest.

---

**Supplemental Figure 1. PAW treatment resulted in variation in primary root length and lateral root density**

Three-day old Arabidopsis seedlings were transferred to N-free media supplemented with diH_2_O (control), 1.75 mM or 2.4 mM NO_3_^-^, with or without H_2_O_2_ for 5 days. Primary root length (A) and lateral root density (B) were quantified in Fiji. A one-way ANOVA with Tukey’s multiple comparison test was performed. N= 40 seedlings.

**Supplemental Figure 2. Principal Component Analysis of root tissue samples.**

Three-day old Arabidopsis seedlings were transferred to N-free media supplemented with PAW4 or 2.4mM NO_3_^-^ as a control. Roots and shoots were harvested one week later, and RNA was extracted and sequenced. A principal component analysis (PCA) was conducted for the sequenced RNA. 4 PAW-treated root replicates (PR) and 3 NO_3_^-^ control-treated root replicates were plotted.

